# Analysis of single-cell RNA-seq data from ovarian cancer samples before and after chemotherapy links stress-related transcriptional profile with chemotherapy resistance

**DOI:** 10.1101/2020.06.06.138362

**Authors:** Kaiyang Zhang, Erdogan Pekcan Erkan, Jun Dai, Noora Andersson, Katja Kaipio, Tarja Lamminen, Naziha Mansuri, Kaisa Huhtinen, Olli Carpén, Johanna Hynninen, Sakari Hietanen, Jaana Oikkonen, Antti Häkkinen, Sampsa Hautaniemi, Anna Vähärautio

## Abstract

Chemotherapy resistance is the greatest contributor to cancer mortality and the most urgent unmet challenge in oncology. In order to reveal transcriptomics changes due to platinum-based chemotherapy we analyzed single-cell RNA-seq data from fresh tissue samples taken at the time of diagnosis and after neoadjuvant chemotherapy from 11 high-grade serous ovarian cancer (HGSOC) patients. With a novel clustering method accounting for patient-specific variability and technical confounders, we identified 12 clusters. Of these, a stress-related transcriptional profile was enriched during chemotherapy and associated significantly to poor progression-free survival (PFS) and disease-free interval (DFI) in deconvoluted bulk RNA-seq data analysis of treatment-naive samples in TCGA cohort. Pan-cancer cell line analysis suggests that patients with high stress-related transcriptional profile may benefit from MEK1/2 inhibitors instead of platinum. Further, high proportion of stromal components and high interaction score between tumor and stromal suggest the tumor cells with high-stress profile actively interact with and potentially recruit stromal cells to their microenvironment already prior to chemotherapy, potentially facilitating protection from chemotherapeutic treatments. In summary, we have identified a stress-related transcriptional profile, which is present at the time of diagnosis, enriched during platinum treatment, independent predictor for poor PFS and DFI, and, based on *in vitro* data, targetable with MEK1/2 inhibitors.

**Translational relevance:** We discovered a stress-related transcriptional profile that is significantly enriched in fresh tissue samples after chemotherapy and is significantly associated with poor progression-free survival in an independent patient cohort. The survival association is independent of age, tumor purity or BRCAness. Therefore, this chemotherapy resistance associated profile is intrinsic and could thus be targeted already in treatment-naive patients. The translation potential of the stress-related transcriptomics profile was further supported by pan-cancer cell line analysis that showed that cell lines with high stress-related transcriptional profile are not affected by platinum, corroborating our results, whereas they were more sensitive to MEK1/2 inhibitors. Taken together, the stress-related transcriptional profile, quantifiable with a set of 35 marker genes, provides a basis for improved prediction of platinum response as well as novel avenues to treat this patient group more effectively.

## Introduction

Chemotherapy resistance is the greatest contributor to cancer mortality and the most urgent unmet challenge in oncology (Holohan et al., 2013; Housman et al., 2014). Platinum-based chemotherapy is prescribed for 10-20% of all cancer patients (NCI, 2014) and presents significant impressive survival advantages, especially in testicular and ovarian cancers (Hanna & Einhorn, 2014; Neijt et al., 1987). Still, the majority of the patients develop resistance to platinum and suffer relapse with limited treatment options. Mechanisms leading to platinum resistance are complex and include platinum uptake and efflux, DNA damage repair, and dysfunctional apoptosis signaling (Galluzzi et al., 2012; Kozłowska et al., 2018). It is, however, poorly understood which of them are active in a patient at relapse, mostly due to paucity of high-quality samples after chemotherapy.

High-grade serous ovarian carcinoma (HGSOC) is the most common and aggressive subtype of ovarian cancer (Koshiyama et al., 2017; Torre et al., 2018). The standard first-line therapy for HGSOC is primary debulking surgery (PDS) followed by platinum-based chemotherapy (Du Bois & Pfisterer, 2005). Platinum-based neoadjuvant chemotherapy (NACT) prior to interval debulking surgery (IDS) is recommended for patients with advanced stage HGSOC when successful PDS is unlikely to be achieved (Wright et al., 2016). Although most HGSOC patients initially respond well to the standard-of-care therapy, the majority develops chemotherapy resistance and relapses leading to a 5-year survival rate of only 43% (Ledermann et al., 2018).

HGSOC is characterized by the universal presence of *TP53* mutations and widespread copy number alterations (CNAs) (Ahmed et al., 2010; Etemadmoghadam et al., 2009). Low frequency of oncogenic mutations and few recurrent CNAs make it difficult to identify targetable genetic lesions (Hoadley et al., 2014; Lheureux et al., 2019). The loss of *TP53* leads to high intratumoral heterogeneity, which is one of the major drivers of therapeutic resistance (Lee et al., 2011; Morris et al., 2016). In addition to genetic heterogeneity, substantial evidence demonstrates that the transcriptional heterogeneity, which is associated with genetic alterations and other sources of variations, such as epigenetic alterations and tumor microenvironment, plays important roles in cancer progression and treatment resistance (Almendro Navarro et al., 2014; González-Silva et al., 2020). Previous studies have revealed transcriptional subtypes for HGSOC (Chen et al., 2015). However, the analyses using bulk transcriptomes can be severely biased by cell type compositions and have limited ability to capture the heterogeneity within a tumor as multiple subtypes can co-exist in the same tumor. A recent work using single-cell RNA sequencing (scRNA-seq) has characterized six cell subtypes of fallopian tube epithelium and identified a epithelial-mesenchymal transition (EMT)-high subtype of HGSOC with poor prognosis (Hu et al., 2020). However, the transcriptional changes due to chemotherapy have remained elusive.

Here, we performed scRNA-seq on fresh tumor samples collected at the time of diagnosis and after NACT from 11 patients to better understand the tumor cell transcriptional profiles in HGSOC and their changes in response to chemotherapy. We identified 12 subpopulations characterized by 10 distinct gene communities. A pre-existing stress associated subpopulation in pre-NACT samples was found to be expanded after chemotherapy. Further validation using bulk RNA-seq data suggests the stress-related profile independently predicts platinum-based chemotherapy resistance and patient prognosis already in treatment-naive samples.

## Results

### Identification of tumor cell clusters in HGSOC

To interrogate the intra-tumor transcriptomic heterogeneity and identify biomarkers related to chemotherapy resistance in HGSOC, we performed scRNA-seq on primary (pre-NACT) - interval (post-NACT) fresh tumor pairs collected from 11 patients (Fig. 1A). Major cell types, i.e., tumor cells, stromal cells and immune cells were identified based on graph-based clustering (Stuart et al., 2019) and acknowledged markers (Fig. S1). As HGSOC is characterized by chromosomal instability and extensive CNAs (Cope et al., 2013; Kuo et al., 2009), we filtered out cells which lack inferred CNAs and have their inferred CNA profiles clustered together with stromal cells. A total of 8,806 tumor cells, with a median of 3677 detected genes per cell, passed our quality controls. The scRNA-seq downstream analysis focused on the tumor cells.

**Figure 1.**
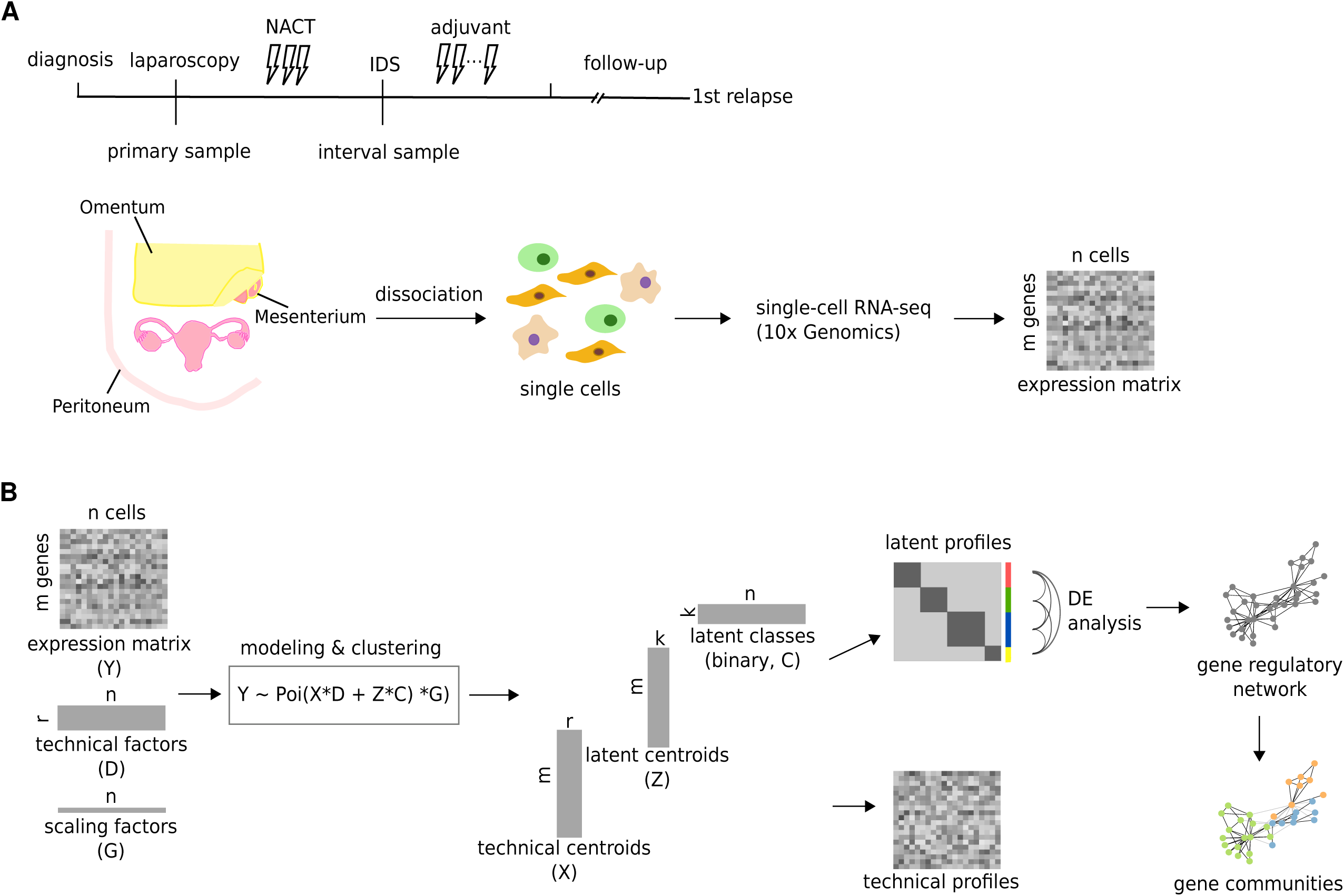
Overall workflow of the sample preparation and data analysis. **A**, Diagram shows the sample collection and processing. We collected fresh tumor samples from 11 high-grade serous ovarian cancer (HGSOC) patients before and after neoadjuvant chemotherapy (NACT). scRNA-seq was performed using the 10x Genomics Chromium platform. **B**, Schematic of our approach to cluster cells and detect co-expressed gene communities. The observed single cell expression profiles (*Y*) is a mixture of latent transcriptional profiles and nuisance contributions. Given the observations *Y*, the known nuisance factors *D*, known size factors *G* and the number of latent clusters *k*, we can estimate the latent nuisance contributions *X*, latent transcriptional profiles *Z* and the latent cluster memberships *C* using an Expectation-Maximization (EM) algorithm.

To identify tumor cell clusters in the presence of unwanted sources of variation, including patient-specific variability and technical confounders, e.g., percentages of unique molecular identifier (UMI) counts originating from mitochondrial genes, we separated the expression readouts into nuisance contribution profiles and latent cluster contribution profiles using an iterative Expectation-Maximization (EM) algorithm with multiple random restarts (Fig. 1B, Methods). First we selected the number of clusters to be 12 based on Bayesian information criterion (BIC, Fig. S2) by performing the procedure on up to 25 clusters using 10 restarts. Second, we performed 200 restarts for the selected number of clusters to ascertain a globally good clustering, and the clusters and profiles computed with the highest likelihood were used for subsequent analysis. Three out of 12 clusters (C3, C9, C10) were patient-specific with over 95% of cells from an individual patient, and the majority of cells in C3 were from a primary tumor while C9 and C10 mainly contained cells from interval tumors. All other nine clusters were shared by primary and interval tumors from multiple patients (Fig. 2A-C). To further evaluate the robustness of the cell clusters, we repeated the analysis with another 200 random restarts and the detected clusters were found to be highly consistent with the presented cell clusters (Fig. S3).

**Figure 2.**
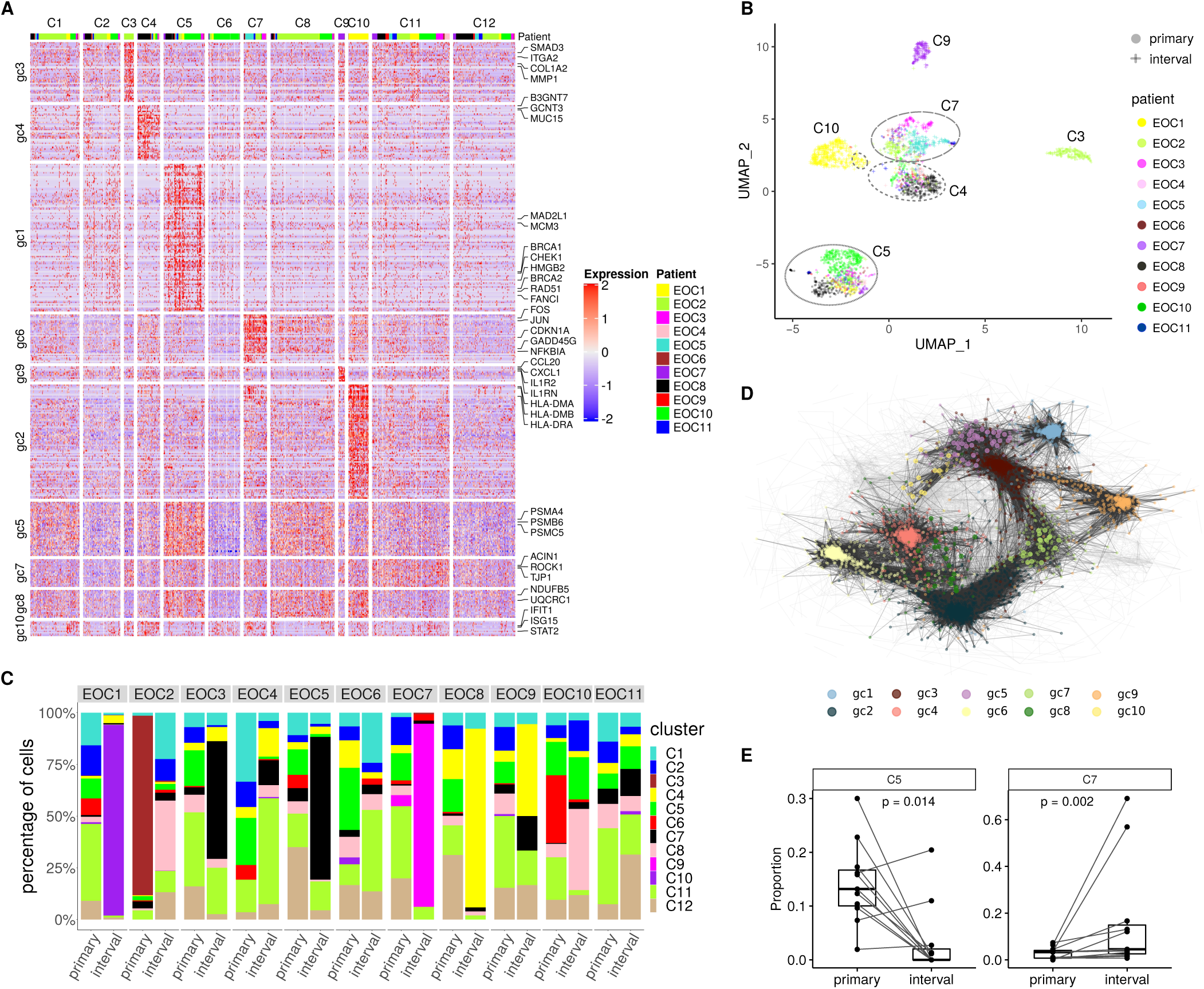
Identification of 10 gene communities characterizing 12 subpopulations of HGSOC tumor cells. **A**, Heatmap of the expression of the 10 distinct gene communities in the 12 identified cell clusters. Rows correspond to genes and columns to cells. The color bar on the top denotes the patients. **B**, The uniform manifold approximation and projection (UMAP) plot shows the clusters within HGSOC tumor cells, colored by the patient. The 4 clusters with no distinguishing gene communities enriched are not shown in this plot. **C**, Stacked bar plot shows the proportion of cells in each identified cluster in each tumor. Colors of the bars denote the clusters. **D**, Force-directed embedding layout of the network with the 4742 differentially expressed genes (DEGs) as vertices and their Pearson correlations of the likelihood-ratio test statistic (LRT) as edges. Colors denote the gene communities to which the cells are assigned, and the unassigned genes are omitted. **E**, Box plots comparing the proportion of cells in cluster C5 (left) and C7 (right) between the primary and interval samples.

### Tumor cell subpopulations are characterized by distinct gene communities

To characterize the identified tumor cell clusters and find co-expressed gene communities, we built a gene network using the 4742 genes that were differentially expressed between at least one pair of the identified cell clusters. The significant Pearson correlations (*ρ* > 0.8, *p* < 0.01) of the likelihood-ratio test statistic (LRT) for each gene between all pairs of clusters were defined as the edges of the network. We detected 916 communities in the network via random walks (Pons & Latapy, 2005), and the 10 communities consisting of more than 30 genes per community were kept for further analysis (Fig. 2D).

For each gene community, only the functionally annotated genes with highest coreness centrality were retained. The relations between the identified cell clusters and the gene communities with the corresponding functional enrichment are shown in Fig. S4 and Table S2. Gene communities gc2, gc3 and gc9 were overrepresented in patient-specific clusters, C10, C3, C9, respectively. gc2 was associated with major histocompatibility complex (MHC) class II antigen presentation genes (e.g., *HLA-DPA1, HLA-DMA, HLA-DOA, HLA-DQA1, HLA-DRA*), gc3 was enriched in TGF-beta signaling (e.g., *SMAD3, ITGA2, COL1A2*) and gc9 was related to cytokines and chemokines (e.g., *IL1R2, CCL20, CXCL1*). Four gene communities (gc5, gc7, gc8, gc10) were shared by multiple cancer cell clusters. They were associated with translation, RNA processing and apoptosis, TCA cycle, and interferon signaling, respectively.

Gene community gc4 was enriched in mucin-type O-linked glycosylation (e.g., *MUC4, MUC5B, MUC15, B3GNT7-8*), which is related to post-translational modifications and protein integrity and has been shown to be involved in tumor growth and progression (Burchell et al., 2018; Gupta et al., 2020).

Gene community gc1 was strongly associated with cell cycle (e.g., *MAD2L1, MCM2-7, CHEK1, HMGB2*) and DNA repair related pathways, including homologous recombination (e.g., *BRCA1/2, RAD51, POLD1*), Fanconi anemia pathway (e.g., *FANCG, FANCI, FANCD2*), base excision repair, nucleotide excision repair and mismatch repair pathways. This gene community was overrepresented in cell cluster C5 (Fig. 2A, Fig. S4), which contained on average 14.3% cells from each primary tumor and 3.3% cells from each interval tumor (Fig. 2E). Lower proportion of cycling cells suggests that platinum-taxane chemotherapy has induced cell cycle arrest in the interval samples.

Gene community gc6, consisting of 35 genes, was highly enriched with early response genes (e.g., *CEBPB, FOS, JUN, DDIT3*) and pro-survival genes (e.g., *GADD45B* and *GADD45G*). Cell cluster C7 was characterized by gc6, and we observed increased proportions of interval tumor cells in C7 compared to primary tumor cells (*p* = 0.002, Fig. 2E). We further defined a stress score as the gene set enrichment (ssGSEA) score (Barbie et al., 2009; Subramanian et al., 2005) for gc6 in individual cells or samples. Eight out 11 patients showed a higher stress score in the interval tumor than in the primary tumor (Fig. 3A). To investigate whether the increase of stress score after chemotherapy is unique to the scRNA-seq data or conserved in HGSOC bulk RNAseq data, we computed the stress score based on their deconvoluted cancer cell specific profile for 23 HGSOC bulk samples including 18 primary-interval pairs and eight primary-relapse pairs. Consistently, the interval tumors showed significantly higher stress scores compared to their paired primary tumors in the bulk samples (*p* = 0.024, Fig. 3B left). Notably, higher stress scores were also observed in relapse tumors (*p* = 0.0078, Fig. 3B right). Even when accounting for anatomical sites, the stress score varied significantly between primary and interval (*p* = 1.45e-07 for partial rank correlation), primary and relapse (*p* = 0.067 for partial rank correlation) tumors. The results from the bulk RNA-seq show that the relevance of the stress-related transcriptional profile in chemotherapy resistance is not an artifact due to single-cell dissociation.

**Figure 3.**
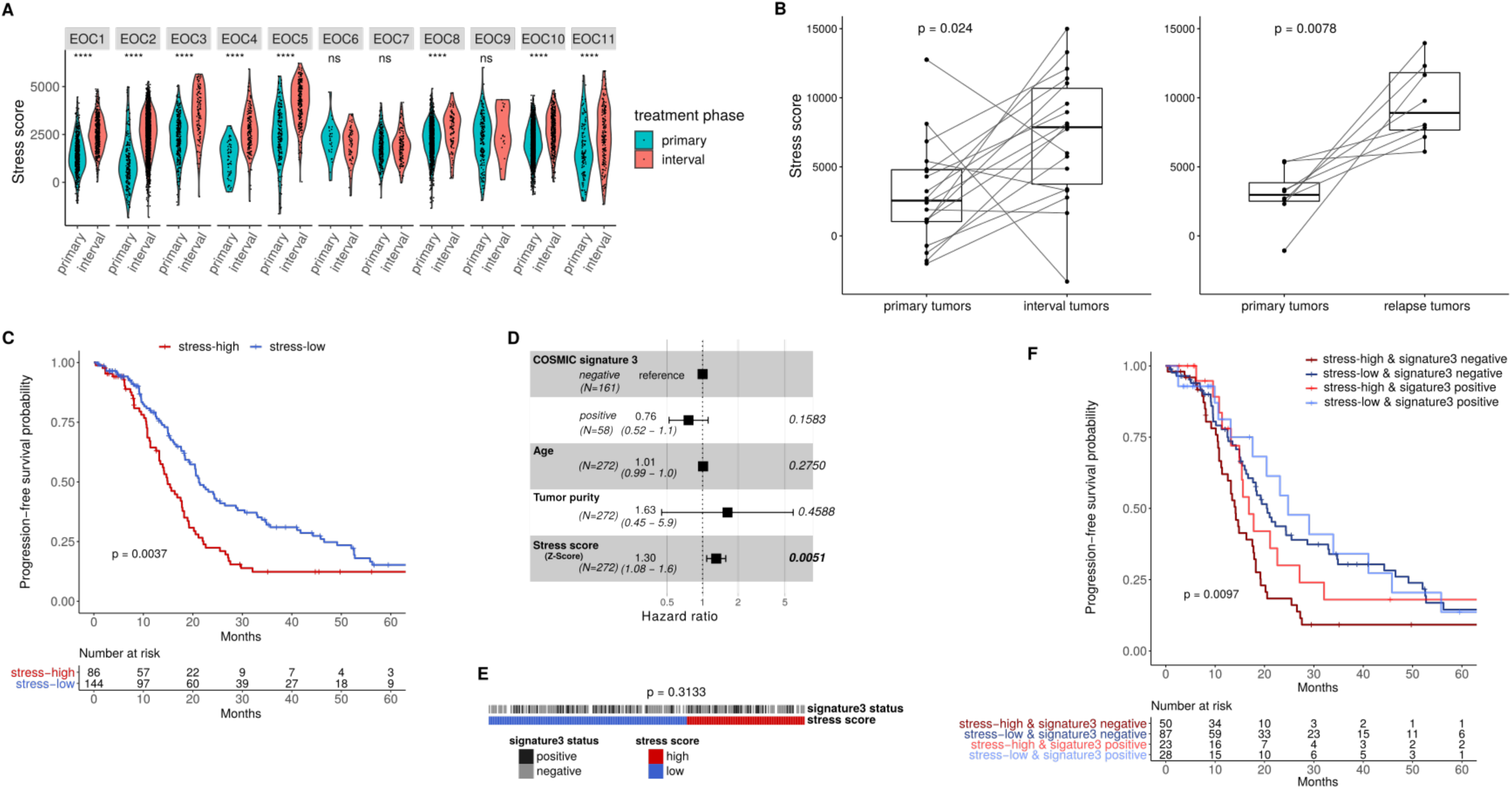
Stress score is associated with chemotherapy response and patient prognosis. **A**, Violin plots comparing the stress score between cells in primary and interval tumors. Violins are colored by the treatment phase and each dot represents a cell. The statistical significance by Wilcoxon rank-sum test is indicated by asterisks (****, *p* < 0.0001; ***, *p* < 0.001; **, *p* < 0.01; *, *p* < 0.05 or ns, *p* > 0.05). **B**, Box plots comparing the stress scores between primary and interval tumors (left, paired Wilcoxon rank-sum test p = 0.024) and primary and relapse tumors (right, paired Wilcoxon rank-sum test p = 0.0078) in the HERCULES cohort. **C**, Kaplan–Meier curves on progression-free survival for stress-high and stress-low patients (log-rank test, *p* = 0.0037) from the TCGA cohort. The number of patients at risk is listed below the survival curves for each time point. **D**, Forest plot showing hazard ratios, their confidence intervals and p-values based on a multivariate Cox proportional hazards regression model studying progression-free survival in relation to COSMIC signature 3 status, age at diagnosis, tumor purity and stress score. Significant p values (P < 0.05) are shown in bold. **E**, COSMIC signature 3 status is not enriched in stress-high or low patients in the TCGA cohort (Fisher exact test, *p* = 0.3133). **E**, Kaplan–Meier curves on progression-free survival for patients from the TCGA cohort with stress-high & signature 3 negative (ii) stress-low & signature 3 negative (iii) stress-high & signature 3 positive and (iv) stress-low & signature 3 positive. The log-rank test p value is 0.0097. The number of patients at risk is listed below the survival curves for each time point.

### The stress-related gene community is associated with poor prognosis

We next turned to the bulk HGSOC RNAseq dataset from The Cancer Genome Atlas (TCGA) (Network & The Cancer Genome Atlas Research Network, 2011) and tested the association between the stress scores and patient outcome. The cell type specific proportion and expression profiles were estimated using PRISM, a latent statistical framework to simultaneously extract the sample composition and cell type specific expression profiles (Häkkinen et al., 2019). After deconvolution, the stress scores were computed using cancer cell specific expression profiles. We identified 86 stress-high tumors that were significantly enriched with the stress gene community gc6 (*p* < 0.05) and 145 stress-low tumors (*p* > 0.5). Patients with stress-high tumors had significantly shorter progression free survival (PFS, *p* = 0.0037 in log-rank test) and disease-free interval (DFI, *p* = 0.015 in log-rank test) than the patients with stress-low tumors (Fig. 3C, Fig. S5).

HGSOC patients with COSMIC signature 3 (BRCA signature) are associated with dysfunctional *BRCA1/2* and better response to chemotherapy (Patch et al., 2015). Therefore, we investigated whether the lack of signature 3 could explain the poor prognosis of the stress-high patients. We found that signature 3 status is not enriched in stress-high or low patients (Fig. 3E, Fisher exact test *p* = 0.3133). Multivariate Cox regression analysis showed that the stress score was significantly associated with PFS (*p* = 0.0051, Fig. 3D) and DFI (*p* = 0.0339, Fig. S5B) independently from the effect of age, tumor purity and signature 3 status. We stratified both stress status and signature 3 status into four groups: (*i*) stress-high & signature 3 negative (*ii*) stress-low & signature 3 negative (*iii*) stress-high & signature 3 positive and (*iv*) stress-low & signature 3 positive. Short PFS and DFI in stress-high patients were observed in both signature3 positive and signature3 negative groups (Fig. 3F). As TCGA has only primary tumors, these results suggest that the stress-related transcriptional profile pre-exists in the untreated tumors. Our results show that the stress score can identify poor-prognostic HGSOC patients independently of the BRCA signature.

### High-stress tumors show an altered microenvironment

Cellular stress has a pivotal role in remodeling the tumor microenvironment (Chang et al., 2017; Koumenis et al., 2013). Accordingly, we tested for an association between the stress scores and the microenvironment compositions in the TCGA cohort. We found significantly higher proportions of stromal (*p* = 2.4e-09) and immune (*p* = 3e-04) components in the stress-high tumors (Fig. 4A).

**Figure 4.**
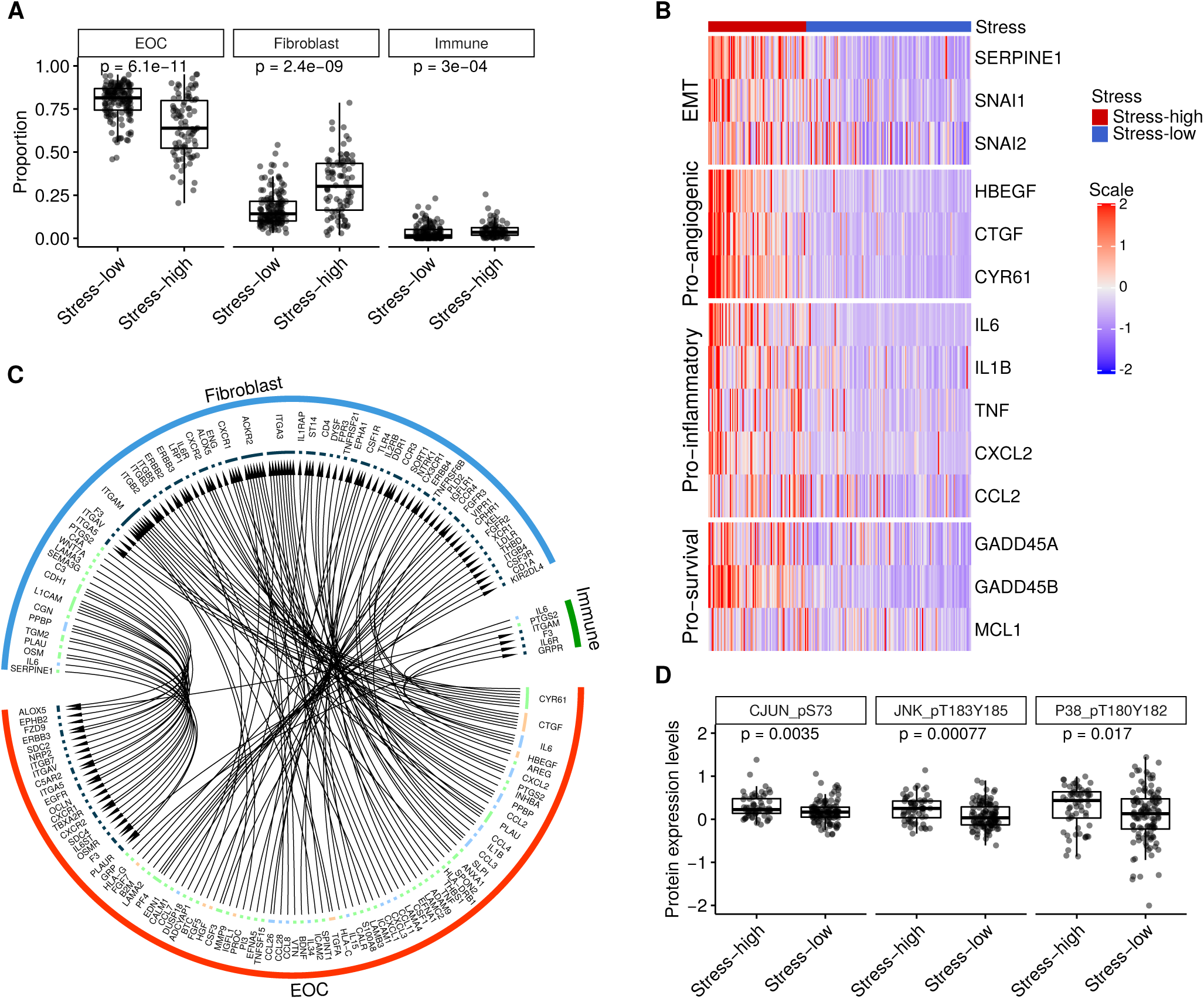
Stress-high tumors show altered microenvironments. **A**, Box plots comparing the proportion of tumor (*p* = 6.1e-11), stromal (*p* = 2.4e-09) and immune (*p* = 3e-04) components between stress-high and stress-low patients from the TCGA cohort. Each dot represents a sample, the p values were computed using Wilcoxon rank-sum test. **B**, Heatmap shows the tumor cell specific expression of epithelial to mesenchymal transition (EMT), pro-angiogenic, pro-inflammatory and pro-survival marker genes in the 86 stress-high and 145 stress-low samples from TCGA cohorts. Samples are sorted by the stress scores. **C**, Chord diagram shows interactions between the tumor component and stromal, immune components that are increased in stress-high tumors compared to stress-low tumors (log_10_FDR < -5 & fold change > 5). **D**, Box plots comparing the protein expression levels of c-JUN_pS73 (*p* = 0.0035), JNK_pT183Y185 (*p* = 0.00077), P38_pT180Y182 (*p* = 0.017) between stress-high and stress-low patients from the TCGA cohort. P values were computed using Wilcoxon rank-sum test.

We further analyzed the cell type specific expression profiles between stress-high and stress-low tumors. Key pro-inflammatory cytokines (e.g., *IL1B, IL6* and *TNF*), genes involved in EMT (e.g., *SERPINE1, SNAI1* and *SNAI2*), pro-angiogenic cytokines (e.g., *HBEGF, CTGF, CYR61*) and pro-survival factors (e.g., *GADD45A, GADD45B, MCL1*) were upregulated in tumor, stromal and immune components of high-stress tumors (Fig. 4B). This suggests that tumor cells with high stress-related profile are more active in recruiting and modulating their surrounding cells to gain survival advantages through cytokine-mediated signaling pathways.

To further characterize the communication between tumor cells and the microenvironment in stress-high tumors, we used an interaction score defined as the product of the ligand expression in one cell type and the corresponding receptor expression in another cell type to quantify the activity of 2,597 previously collected ligand-receptor interactions (Kumar et al., 2018; Wang et al., 2019). The interactions with the highest increase in stress-high tumors were related to the pro-inflammatory, EMT and pro-survival cytokines. We also observed significantly higher scores for interactions related to laminins (*LAMA2, LAMA4, LAMB3* and *LAMC2*) that bind to integrin receptors and collagens (*COL5A2, COL11A1*) binding to *DDR1* in stress-high tumors (Fig. 4C).

To confirm the occurrence of the stress-related profile at the protein level, we analyzed the TCGA HGSOC reverse-phase protein array (RPPA) data, which is available for 224 proteins in 60 stress-high and 115 stress-low tumors. The phosphorylated c-Jun (CJUN_pS73, *p* = 0.0035) and its upstream kinase phospho-JNK (JNK_pT183Y185, *p* = 0.00077) were increased in the stress-high tumors (Fig. 4D). Moreover, we observed higher levels of phosphorylated p38-α (P38_pT180Y182, *p* = 0.017, Fig. 4D), supporting the induction of JNK and p38 MAPK signaling pathways also on the protein level in stress-high tumors.

### Identification of drugs potentially targeting the stress-high tumors

To identify drugs that can effectively target the stress-high tumor cells *ex vivo*, we obtained the gene expression and GDSC2 drug sensitivity datasets from the Genomics of Drug Sensitivity in Cancer (GDSC) (Yang et al., 2013). Among the 1,018 cell lines with gene expression data available, we identified 233 stress-high cell lines spanning various cancer types (Fig. 5A).

**Figure 5.**
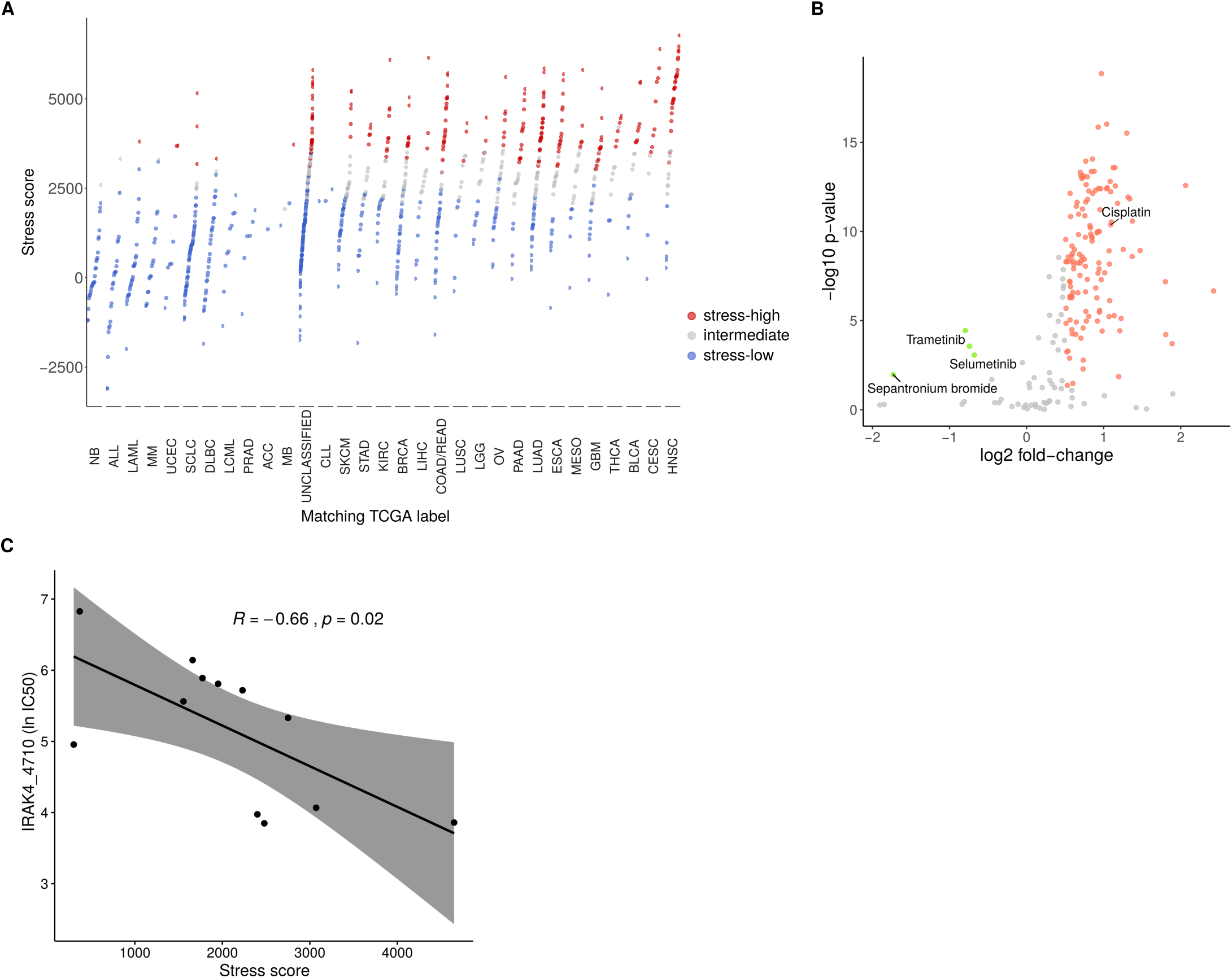
Association between stress scores and drug sensitivity in the GDSC2 cell line dataset. **A**, Dot plot shows the stress-high (233) and stress-low (504) cells lines in the GDSC2 dataset. Colors denote the stress status. **B**, Volcano plot shows the compounds with significantly different half-maximal inhibitory concentration (IC50) between stress-high and stress-low cell lines. The red and green dots correspond to the compounds with significantly different IC50 (log_2_fold-change > 0.5, Wilcoxon rank-sum test *p* <0.05). **C**, Scatter plot shows the correlation between the stress scores and lnIC50 across the 12 ovarian cancer cell lines harboring *TP53* mutations (*R* = -0.66, *p* = 0.02).

We compared the drug response for 192 compounds between stress-high and stress-low cell lines, consistent with our analysis of patient samples (Fig. 3), significantly increased half-maximal inhibitory concentration (IC50) of cisplatin were observed in stress-high cell lines (log2 fold-change = 1.09, *p* = 4.19e-11, Fig. 5B). Notably, the stress-high cell lines were more sensitive to MEK1/2 inhibitors, Trametinib (log2 fold-change = -0.79, *p* = 3.63e-05, Fig. 5B) and Selumetinib (log2 fold-change = -0.68, *p* = 8.49e-04, Fig. 5B) with significantly lower IC50.

Considering the cancer type specific variability, we next quantified the correlation between stress scores and lnIC50 across the 12 ovarian cancer cell lines harboring *TP53* mutations. In this relatively small subset, we found a significant negative correlation between the stress score and the lnIC50 of the IRAK4 inhibitor (IRAK4_4710, R = -0.66, *p* = 0.02, Fig. 5C). IRAK4 has been reported to play an important role in IL1-induced JNK and p38 activation (Rhyasen & Starczynowski, 2015; Suzuki et al., 2002).

## Discussion

We analyzed fresh tissue samples taken before and after chemotherapy from 11 HGSOC patients and subjected to scRNA-seq to identify molecular processes that drive resistance to platinum-taxane combination therapy. The scRNA-seq results were validated with and expanded to bulk RNA-seq data from 23 HGSOC patients as well as with 272 HGSOC patients available in TCGA.

By using a novel clustering method accounting for patient-specific variability and technical confounders, we identified 10 distinct transcriptional profiles, of which seven were shared across different patients. Our results reveal a stress-related transcriptional profile, which is induced by chemotherapy and related to poor prognosis in treatment-naive patients. The identified stress-related transcriptional profile can be characterized by a set of only 35 marker genes, providing a basis for identification of poor-prognostic patients as well as developing new treatment strategies. Importantly, our findings imply that the expression pattern is associated with both intrinsic and adaptive resistance, and could be targeted in both treatment-naive patients predicted to have poor response, and following chemotherapy to minimize residual disease. Analysis of GDSC2 drug sensitivity data sets suggests IRAK4 or MEK1/2 inhibitors as potential drugs to target the cisplatin resistant stress-high subpopulations in tumors. These results open interesting avenues for mechanistic studies.

Increasing evidence shows that stress inducible genes and pathways can be activated in response to the selective microenvironment forces, such as hypoxia, nutrient deprivation and DNA damage, to help cancer cell survival and progression (Avril et al., 2017; Tsuruo et al., 2003). Upregulation of stress induced immediate early genes (IEGs) *FOS* and *ATF3* has been reported to be associated with increasing resistance to chemotherapeutic agents (Chang et al., 2017; Mukhopadhyay et al., 2001). Baron and colleagues showed that the stress-like cancer cells confer resistance to MEK/BRAF inhibitors in melanoma (Baron et al., 2020). However, most of the studies are based on laboratory models instead of human tumors. The stressed-related profile has been found in other single cell studies in muscle stem cells (van den Brink et al., 2017) and HGSOC (Hu et al., 2020). Hu et al. suggest several genes belonging to the stress-related transcriptional profile to be artifacts due to sample preparation for scRNA-seq experiments. We tested the presence of the stress-related transcriptional profile in bulk RNA-seq samples from HGSOC patients’ fresh tumors without single-cell dissociation in sample handling and were able to identify them as predicted from scRNA-seq data. This suggests that, at least in our data set, the stress-related transcriptional profile is not a dissociation artifact.

Tumors with high stress scores were found to have higher proportions of stromal and immune components and importantly, the association between stress score to PFS and DFI was independent of tumor purity. Ligand-receptor analysis showed the genes enriched in high-stress tumors contained a high number of ligands, and to lesser extent receptors, for the tumor microenvironment interactions to mainly stromal cells. This suggests that tumor cells with high stress-related profile actively interact with especially stromal cells and potentially recruit them to their microenvironment already prior to chemotherapy, thus facilitating protection from chemotherapeutic treatments.

## Materials and Methods

### Human subjects

All patients participating in the study provided written informed consent. The study and the use of all clinical material have been approved by The Ethics Committee of the Hospital District of Southwest Finland (ETMK) under decision number EMTK: 145/1801/2015.

### scRNA-seq sample preparation

Fresh HGSOC tumor specimens were collected from 11 patients at the time of laparoscopy and interval debulking surgery (IDS). Detailed clinical information is shown in Table 1. Immediately after surgery, the specimens were incubated overnight in a mixture of collagenase and hyaluronidase (Department of Pathology, University of Turku) to obtain single cell suspensions, that were then processed with the standard Chromium Single Cell 3′ Reagent Kit v. 2.0 (10x Genomics) protocol for scRNA-seq with Illumina HiSeq4000 (Jussi Taipale Lab, Karolinska Institute and Institute for Molecular Medicine Finland).

**Table 1.** Patient and sample information.

| Patient ID | Age<sup>1</sup> | Treatment | Stage<sup>2</sup> | PFI (days) | Anatomical locations <sup>3</sup> |  |  |  |  |
| --- | --- | --- | --- | --- | --- | --- | --- | --- | --- |
|  |  |  |  |  | scRNA-seq Primary | scRNA-seq Interval | Bulk RNA-seq Primary | Bulk RNA-seq Interval | Bulk RNA-seq Relapse |
| EOC1 | 73 | NACT | IVA | 65 | Peritoneum | Tumor | NA | NA | NA |
| EOC2 | 64 | NACT | IVA | 520 | Mesentery | Omentum | NA | NA | NA |
| EOC3 | 78 | NACT | IVA | 393 | Omentum | Omentum | NA | NA | NA |
| EOC4 | 74 | NACT | IVA | 230 | Omentum | Omentum | NA | NA | NA |
| EOC5 | 67 | NACT | IVA | 14 | Peritoneum | Omentum | Peritoneum | Omentum | NA |
| EOC6 | 67 | NACT | IVB | 36 | Peritoneum | Omentum | NA | NA | NA |
| EOC7 | 68 | NACT | IIIC | 460 | Peritoneum | Peritoneum | Peritoneum | Peritoneum | NA |
| EOC8 | 54 | NACT | IVA | 177 | Omentum | Omentum | Omentum | Omentum | NA |
| EOC9 | 62 | NACT | IIIC | 126 | Omentum | Omentum | NA | NA | NA |
| EOC10 | 72 | NACT | IVA | 83 | Peritoneum | Omentum | NA | NA | NA |
| EOC11 | 62 | NACT | IIIC | 30 | Peritoneum | Omentum | Omentum | Omentum | NA |
| EOC12 | 75 | NACT | IIIC | 210 | NA | NA | Omentum | Mesentery | NA |
| EOC13 | 68 | PDS | IVB | 648 | NA | NA | Omentum | NA | Mesentery |
| EOC14 | 68 | NACT | IIIC | 203 | NA | NA | Omentum | Omentum | NA |
| EOC15 | 71 | NACT | IIIC | 974 | NA | NA | Omentum | Omentum | NA |
| EOC16 | 72 | NACT | IIIC | 0 | NA | NA | Omentum | Omentum | Ascites |
| EOC17 | 67 | NACT | IIIC | 280 | NA | NA | Omentum | Omentum | NA |
| EOC18 | 81 | NACT | IIIC | 721* | NA | NA | Omentum | Ovary | NA |
| EOC19 | 57 | NACT | IVA | 210 | NA | NA | Mesentery | Omentum | NA |
| EOC20 | 71 | NACT | IVB | 27 | NA | NA | Peritoneum | Tumor | NA |
| EOC21 | 77 | NACT | IVB | 511 | NA | NA | Peritoneum | Omentum | NA |
| EOC22 | 68 | NACT | IIIC | 81 | NA | NA | Peritoneum | Peritoneum | Ascites |
| EOC23 | 74 | NACT | IIIC | 91 | NA | NA | Adnexa | Ascites | NA |
| EOC24 | 71 | NACT | IIIC | 183 | NA | NA | Omentum | NA | Ascites |
| EOC25 | 74 | NACT | IIIC | 632 | NA | NA | Peritoneum | Mesentery | NA |
| EOC26 | 62 | NACT | IIIC | 285 | NA | NA | Peritoneum | Omentum | Peritoneum |
| EOC27 | 66 | NACT | IVB | 661 | NA | NA | Peritoneum | Omentum | NA |
| EOC28 | 75 | NACT | IIIC | 19 | NA | NA | Omentum | NA | Ascites |
| EOC29 | 66 | NACT | IIIC | 221 | NA | NA | Peritoneum | NA | Ascites |
| EOC30 | 64 | NACT | IIIC | 174 | NA | NA | Peritoneum | NA | Ascites |

<sup>1</sup> Age at diagnosis, years
<sup>2</sup> Tumor staging was done according to the International Federation of Gynecology and Obstetrics (FIGO) 2014 guidelines
<sup>3</sup> Anatomical locations from which the samples were collected for analyses.
\* No progression, PFI at outcome update
Abbreviations: NACT: Neoadjuvant chemotherapy; PDS: Primary debulking surgery; PFI: Platinum-free interval, days; NA: Not available

### Preprocessing scRNA-seq data

The Cell Ranger Software Suite (Version 3.1.0) was used to perform sample de-multiplexing, alignment, barcode processing and UMI quantification. The reference index was built upon the GRCh38.d1.vd1 reference genome with GENCODE v25 annotation. Shared nearest neighbor (SNN) modularity optimization based clustering from Seurat v3 (Stuart et al., 2019) were used for initial clustering. Three major cell types were revealed based on acknowledged markers: epithelial tumor cells (*WFDC2, PAX8, EPCAM*), stromal cells (*COL1A2, FGFR1, DCN*), and immune cells (*CD79A, FCER1G, PTPRC*). Cells expressing any combinations of *PAX8, DCN* and *PTPRC* were excluded to remove potential doublets. We focused on tumor cells in the subsequent analysis.

The quality measures of each tumor cell were quantified using Seurat. We estimated the cutoffs for each quality measure based on its bimodal distribution and then used four criteria for quality control: (1) the number of reads above 8,192; (2) the number of UMI counts above 4,705; (3) the number of detected genes above 1,552; (4) the percentage of UMI counts originating from mitochondrial genes below 12. Next, we filtered out tumor cells that have their CNA profiles clustered together with stromal cells. The CNAs were inferred using inferCNV with default settings (inferCNV of the Trinity CTAT Project. https://github.com/broadinstitute/inferCNV). We used an agglomerative hierarchical clustering algorithm to cluster the inferred CNA profiles and the tree was cut based on Bayesian Information Criterion (Häkkinen et al., 2019).

### Modeling and clustering scRNA-seq data

We modeled the observed single cell expression profiles as a mixture of latent transcriptional profiles and nuisance expression profiles following a Poisson distribution: 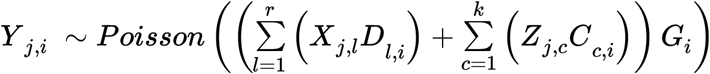, where *l* = 1, 2, *…, r* runs over *r* nuisance factors, *c* = 1, 2, *…, k* runs over *k* latent transcriptional clusters. *Y* _*j,i*_ denotes the observed UMI counts of gene *j* in the *i*^*th*^ cell, *X*_*j,l*_ denotes the expression profile centroid of gene *j* specific to nuisance factor *l, D*_*l,i*_ denotes the design coefficient of the *l*^*th*^ nuisance factor in the *i*^*th*^ cell, *Z*_*j,c*_ denotes the cluster *c* expression profile centroid at gene *j, C*_*c,i*_ ∈ {0, 1} is an indicator of whether the *i*^*th*^ cell belongs to the cluster *c*, and *G*_*i*_ is a cell-specific scaling factor.

Given the observations *Y* _*j,i*_, known nuisances *D*_*l,i*_, known scaling factors *G*_*i*_ and the number of latent clusters *k*, we can estimate the latent nuisance expression centroids *X*_*j,l*_, latent expression centroids *Z*_*j,c*_ and the latent cluster memberships *C*_*c,i*_ using an Expectation-Maximization (EM) algorithm (Dempster et al., 1977). The EM algorithm is constructed on the latent variables 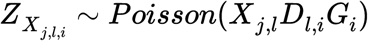 and 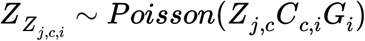, which are the nuisance and cleaned contributions to the expression, respectively. The parameter set *θ* = (*X*_*j,l*_, *Z*_*j,c*_, *C*_*c,i*_) was estimated in two stages: first, the expression centroids *X*_*j,l*_ and *Z*_*j,c*_ can be estimated given *Y* _*j,i*_, *D*_*l,i*_, *C*_*c,i*_ and *G*_*i*_ ; second, the cluster membership *C*_*c,i*_ can be updated given *Y* _*j,i*_, *X*_*j,l*_, *D*_*l,i*_, *Z*_*j,c*_ and *G*_*i*_ (see Supplementary material for the details).

To select the optimal *k*, we fitted our model for *k* = 1, 2, *…,* 25 with 10 different random initial parameter sets for each *k*, and *k* = 12 was selected based on Bayesian Information Criterion (BIC, Fig. S2). We then ran the EM procedure with 200 random initializations for *k* = 12, the maximum likelihood estimates of *X*_*j,l*_, *Z*_*j,c*_ and the resulting *C*_*c,i*_ were used for downstream analysis.

### Differential expression (DE) analysis for scRNA-seq data

Let *Y* _*j,i*_ denote the observed UMI count of gene *j* in the *i*^*th*^ cell, 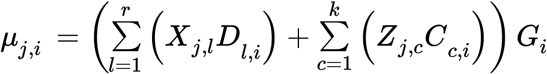 denote the predicted mean expression rate of gene *j* for the *i*^*th*^ cell based on the estimated model parameters *X*_*j,l*_, *Z*_*j,c*_, *C*_*c,i*_. The differential expression between group *g*_1_ and group *g*_2_ for the gene *j* can be assessed by testing the alternative hypothesis 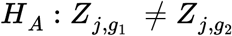 against the null hypothesis 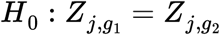. For the former, the likelihood is that attained at the maximum-likelihood estimate (MLE) 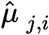,while for the latter, the model is refitted giving 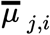, the MLE under *H*_0_. The logarithmic likelihood-ratio test statistic (LRT) for gene *j* is

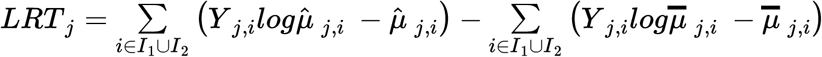

Where *I*_1_ and *I*_2_ denote the indices for the samples in *g*_1_ and *g*_2_, respectively. A p-value for the gene *j* to be differentially expressed between group *g*_1_ and group *g*_2_ can be computed as the probability to the right of the — 2*LRT*_*j*_ for the chi-squared distribution with degrees of freedom equals to the difference in number of parameters, i.e. 1.

### Identification of co-expressed gene communities

Co-expressed gene communities were identified as follows: (1) we conducted DE analysis between each pair of cell clusters, resulting in 66 comparisons. The top 1000 high LRT genes with FDR < 0.01 were selected in each comparison, and this resulted in a total of 4742 genes. (2) the Pearson correlations between these 4742 genes were computed using the LRTs from all 66 comparisons. Correlations with *ρ* > 0.8 and p value < 0.01 were used to build a gene network. (3) we detected 916 communities in the network via random walks with step equals to 3 (Pons & Latapy, 2005), and the 10 communities consisting more than 30 genes were retained. (4) let *V* be the genes in a community, *c*_*j*_ be the coreness of gene *j* and *n*_*max*_ be the number of genes with the maximum coreness (*max*_*j*∈*V*_*c*_*j*_, degeneracy) in that community. If *n*_*max*_*>* 30, then the genes with *c* = *max*_*j*∈*V*_*c*_*j*_ were retained, otherwise, we retained the top 30 genes ranked by coreness. (5) gene set over-representation analysis was performed for the remaining genes in each community using the ConsensusPathDB (Kamburov et al., 2013). We further reduced the redundancy of each gene community with number of genes above 20 by applying the following filters: (*i*) only genes overlapped with significantly overrepresented gene sets (FDR < 0.05, size < 500) were kept; (*ii*) biclustering was applied to the binary matrix of the presence/absence of each gene in each significantly overrepresented gene set using the R package blockcluster (Bhatia et al., 2017), and the gene clusters that have less than 3% presence in any of the gene-set clusters were excluded (Fig. S6). After filtering, the number of genes per community varied between 11 and 106. The genes in each community are listed in Table S1.

### Defining stress scores

We defined the stress score as the gene set enrichment score of our identified stress-associated gene community (gc6) in individual cells and samples, which was computed using ssGSEA (Barbie et al., 2009; Subramanian et al., 2005). Samples with permutation test p-value below 0.05 were considered stress-high while samples with p-value above 0.5 were considered stress-low.

### Bulk tumor expression data

We acquired 18 primary-interval sample pairs and eight primary-relapse sample pairs from 23 patients in the HERCULES cohort (http://www.project-hercules.eu/). The sample collection, data quality control, alignment, quantification were performed as we have previously described (Häkkinen et al., 2019). Briefly, read pairs were trimmed using Trimmomatic (Version 0.33, (Bolger et al., 2014)) and aligned to the GRCh38.d1.vd1 reference genome with GENCODE v25 annotation using STAR (Version 2.5.2b) (Dobin et al., 2013). Gene level effective counts were quantified using eXpress (Version 1.5.1-linux x86 64) (Roberts & Pachter, 2013).

TCGA RNA-Seq data of ovarian serous cystadenocarcinoma (OV, illuminahiseq_rnaseqv2-RSEM_genes_normalized) was downloaded from the Broad Firehose (https://gdac.broadinstitute.org/), along with the clinical annotations. The primary tumors from 272 patients with advanced HGSOC (grade: G2-4, stage: IIIA-IV) were included in our analysis.

The proportions of tumor, stromal and immune components and the cell type specific expression profiles for HERCULES and TCGA samples were estimated using PRISM (Häkkinen et al., 2019).

### TCGA reverse phase protein array (RPPA) data

The RPPA data (replicates-based normalization) for TCGA ovarian serous cystadenocarcinoma samples (TCGA-OV-L4) was downloaded from the Cancer Proteomics Atlas (TCPA, https://www.tcpaportal.org/tcpa/download.html).

### Cell line expression and drug sensitivity data

The GDSC2 drug sensitivity data and the Robust Multichip Average (RMA) normalized expression data were downloaded from the Genomics of Drug Sensitivity in Cancer (GDSC) ftp server (ftp.sanger.ac.uk/pub4/cancerrxgene/releases/).

### Data and code availability

Raw scRNA-Seq expression data will be available at The European Genome-phenome Archive (EGA). The codes for analyses are available at GitHub (https://github.com/KaiyangZ/scRNAseq-HGSOC).

## Acknowledgements

This work is supported financially by the European Union’s Horizon 2020 research and innovation programme under Grant Agreement No. 667403 for HERCULES (Comprehensive Characterization and Effective Combinatorial Targeting of High-Grade Serous Ovarian Cancer via Single-Cell Analysis), the Academy of Finland (Projects No. 292402, 325956, 314395, 289059, 322927, 294023 and 319243), the Sigrid Jusélius Foundation, and the Cancer Foundation Finland. We thank CSC-IT Center for Science Ltd. for compute resources. The results published here are in part based upon data generated by TCGA managed by the NCI and NHGRI. Information about TCGA can be found at https://cancergenome.nih.gov/. Single-cell RNA sequencing was performed with Illumina HiSeq4000 at Jussi Taipale Lab, Karolinska Institute and Institute for Molecular Medicine Finland.

