## Supplementary material for "Analysis of single-cell RNA-seq data from ovarian cancer samples before and after chemotherapy links stress-related transcriptional profile with chemotherapy resistance"

### Modeling and clustering sc-RNAseq data

**Model structure.** We assume each observed single cell expression profiles is a mixture of a latent cellular states and nuisance expression profiles, and can be approximated by a Poisson distribution:

$$Y_{j,i} \sim Poisson\left(\left(\sum_{l=1}^{r}\left(X_{j,l}D_{l,i}\right)+\sum_{c=1}^{k}\left(Z_{j,c}C_{c,i}\right)\right)G_i\right) \quad (1)$$

where $Y_{j,i}$ denotes the observed UMI counts of gene $j$ in the $i^{th}$ cell, $X_{j,l}$ denotes the expression profile centroid of gene $j$ specific to nuisance factor $l$, $D_{l,i}$ denotes the design coefficient of the $l^{th}$ nuisance factor in the $i^{th}$ cell, $Z_{j,c}$ denotes the cluster $c$ expression profile centroid at gene $j$, $C_{c,i} \in \{0,1\}$ is an indicator of whether the $i^{th}$ cell belongs to the cluster $c$, and $G_i$ is a cell-specific scaling factor.

**Parameter estimation.** Given $Y_{j,i}$, $D_{l,i}$, $G_i$ and the number of latent clusters $k$, we can estimate $X_{j,l}$, $Z_{j,c}$ and $C_{c,i}$ using an Expectation-Maximization (EM) algorithm (Dempster, Laird, and Rubin 1977), which is constructed on the latent variables $Z_{X_{j,l,i}}$, $Z_{Z_{j,c,i}}$ and the observations $Y_{j,i}$:

$$\begin{aligned} Z_{X_{j,l,i}} &\sim Poisson(X_{j,l}D_{l,i}G_i) \\ Z_{Z_{j,c,i}} &\sim Poisson(Z_{j,c}C_{c,i}G_i) \\ Y_{j,i} &= \sum_{l=1}^{r} Z_{X_{j,l,i}} + \sum_{c=1}^{k} Z_{Z_{j,c,i}} \end{aligned} \quad (2)$$

The EM iteration alternates between an E-step, which evaluates the expected likelihood over $Z_{X_{j,l,i}}$ and $Z_{Z_{j,c,i}}$ using the current set of parameters and a M-step, where the parameters are updated by maximizing the expected likelihood obtained in the E-step.

The parameter set $\theta = (X_{j,l}, Z_{j,c}, C_{c,i})$ is estimated in two stages: first, we estimate the expression centroids $X_{j,l}$ and $Z_{j,c}$ given $Y_{j,i}$, $D_{l,i}$, $C_{c,i}$ and $G_i$; second, the cluster membership $C_{c,i}$ is updated given $Y_{j,i}$, $X_{j,l}$, $D_{l,i}$, $Z_{j,c}$ and $G_i$.

In the first stage, the parameter estimation algorithm is as described in (Häkkinen et al. 2019). The full model likelihood from Eq (1) is given by:

$$logL(\mathbf{Y}) = \sum_{j=1}^{m}\sum_{i=1}^{n}\left(Y_{j,i}log\left(\sum_{l=1}^{r}X_{j,l}D_{l,i}G_i+\sum_{c=1}^{k}Z_{j,c}C_{c,i}G_i\right)-\left(\sum_{l=1}^{r}X_{j,l}D_{l,i}G_i+\sum_{c=1}^{k}Z_{j,c}C_{c,i}G_i\right)\right)$$

(3)

While the latent model likelihood is given by

$$logL(\mathbf{Z}) = \sum_{j=1}^{m}\sum_{i=1}^{n}\sum_{l=1}^{r}Z_{X_{j,l,i}}logX_{j,l}D_{l,i}G_i-X_{j,l}D_{l,i}G_i+\sum_{j=1}^{m}\sum_{i=1}^{n}\sum_{c=1}^{k}Z_{Z_{j,c,i}}logZ_{j,c}C_{c,i}G_i-Z_{j,c}C_{c,i}G_i$$

(4)

Let $\theta^0=(X_{j,l}^0,\ Z_{j,c}^0,\ C_{c,i}^0)$ be the current set of parameters and

$$E_{Z_{X_{j,l,i}}}=E\left[Z_{X_{j,l,i}}|\theta^0,\ \sum_{l=1}^{r}Z_{X_{j,l,i}}+\sum_{c=1}^{k}Z_{Z_{j,c,i}}=Y_{j,i}\right]$$

$$E_{Z_{Z_{j,c,i}}}=E\left[Z_{Z_{j,c,i}}|\theta^0,\ \sum_{c=1}^{r}Z_{X_{j,l,i}}+\sum_{c=1}^{k}Z_{Z_{j,c,i}}=Y_{j,i}\right]$$

The derivatives of $logL(\mathbf{Z})$ with respect to $X_{j,l}$ and $Z_{j,c}$ are given by:

$$\frac{\partial logL(\mathbf{Z})}{\partial X_{j,l}}=\sum_{i=1}^{n}\frac{E_{Z_{X_{j,l,i}}}}{X_{j,l}}-D_{l,i}G_i$$

(5)

$$\frac{\partial logL(\mathbf{Z})}{\partial Z_{j,c}}=\sum_{i=1}^{n}\frac{E_{Z_{Z_{j,c,i}}}}{Z_{j,c}}-C_{c,i}G_i$$

and $logL(\mathbf{Z})$ is maximized at:

$$\widehat{X}_{j,l}=\frac{\sum_{i=1}^{n}E_{Z_{X_{j,l,i}}}}{\sum_{i=1}^{n}D_{l,i}G_i}$$

(6)

$$\widehat{Z}_{j,c} = \frac{\sum_{i=1}^{n} E_{Z_{Z_{j,c,i}}}}{\sum_{i=1}^{n} C_{c,i} G_i}$$

In the second stage, given $Y_{j,i}, D_{l,i}$ , $G_i$ and updated $X_{j,l}$ , $Z_{j,c}$ in the first stage, we update $C_{c,i}$ by:

$$\widehat{C}_{c,i} = \begin{cases} 1, \text{ iff c= arg} \max_{c=1,2,\ldots,k} \sum_{i=1}^{n} E_{Z_{Z_{j,c,i}}} log Z_{j,c} C_{c,i} G_i - Z_{j,c} C_{c,i} G_i \\ 0, \text{ otherwise} \end{cases} \tag{7}$$

To obtain the latent mean of $Z_{Z_{j,c,i}}$ and $Z_{X_{j,c,i}}$ corresponding to the updated parameters, consider:

$$\begin{aligned} Z_{X_{j,l,i}} &\sim Poisson(X_{j,l} D_{l,i} G_i) \\ Z_{Z_{j,c,i}} &\sim Poisson(Z_{j,c} C_{c,i} G_i) \end{aligned} \tag{8}$$

$$Z_{X_{j,l,i}} + Z_{Z_{j,c,i}} | \sum_{l=1}^{r} Z_{X_{j,l,i}} + \sum_{c=1}^{k} Z_{Z_{j,c,i}} = Y_{j,i} \sim \mathcal{B}(Y_{j,i}, \frac{Z_{X_{j,l,i}} + Z_{Z_{j,c,i}}}{\sum_{l=1}^{r} Z_{X_{j,l,i}} + \sum_{c=1}^{k} Z_{Z_{j,c,i}}})$$

For $K \sim \mathcal{B}(y, p)$ :

$$E[K|y] = py \tag{9}$$

Finally, we summarize the parameter estimation procedure as follows:

1. Choose initial values for the parameters $\theta^0 = (X^0_{j,l},\ Z^0_{j,c},\ C^0_{c,i})$ . We initialize $X_{j,l} = 1$ , $Z_{j,c} = 1$ and $C_{c,i}$ uniform random.
2. Obtain the latent mean of $Z_{Z_{j,c,i}}$ and $Z_{X_{j,c,i}}$ using Eq. (8) and (9).
3. Update $X_{j,l}$ , $Z_{j,c}$ using Eq. (6).
4. Obtain the latent mean of $Z_{Z_{j,c,i}}$ and $Z_{X_{j,c,i}}$ using Eq. (8) and (9).
5. Update $C_{c,i}$ using Eq.(7).
6. Repeat 2-5 until convergence.

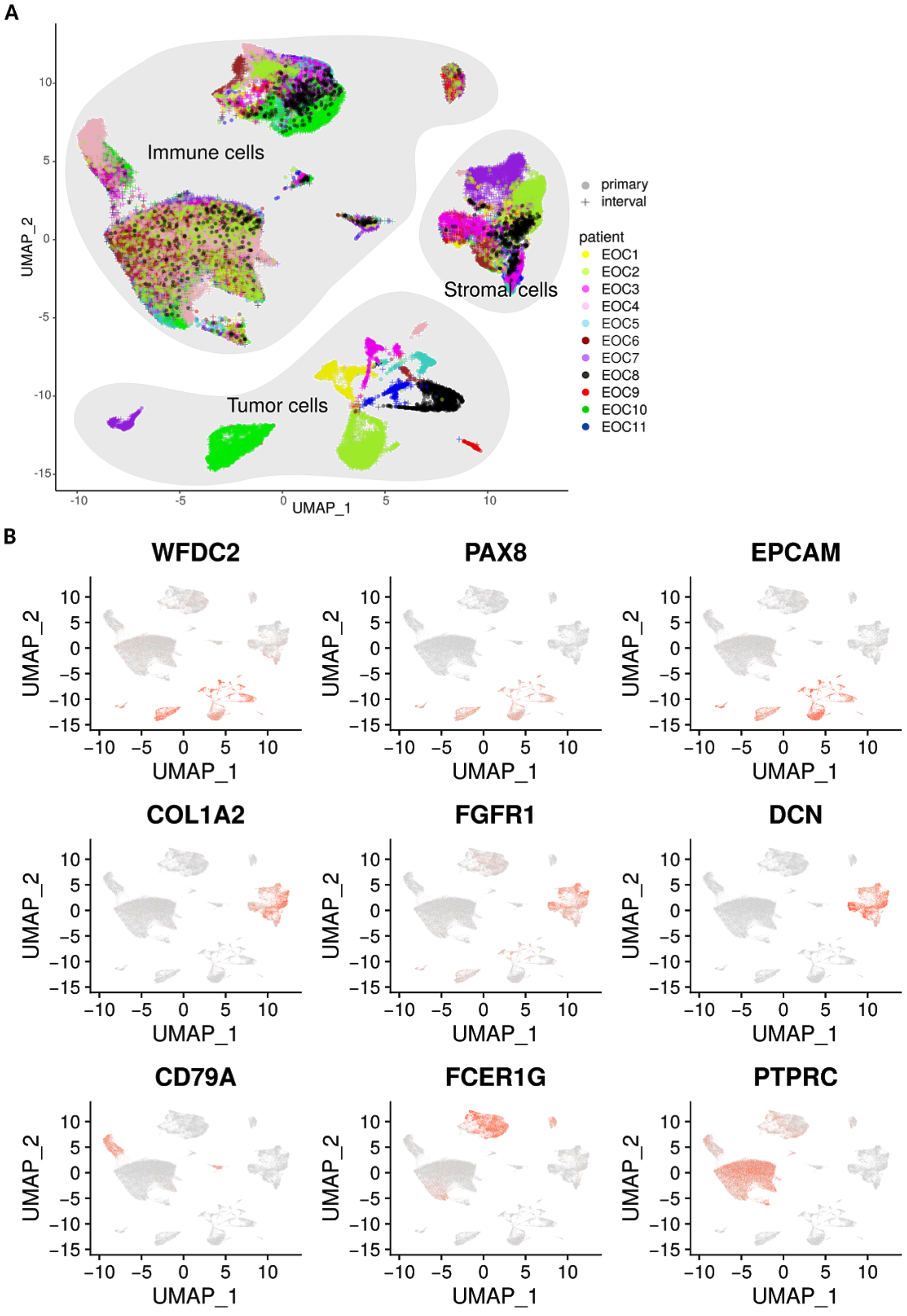


**Supplementary Figure S1**. **Identification of tumor, stromal and immune cell populations from 11 high-grade serous ovarian cancer (HGSOC) patients before and after neoadjuvant chemotherapy (NACT). A,** The uniform manifold approximation and projection (UMAP) plot shows tumor, stromal and immune cell populations, colored by patient and the shapes indicate treatment phase. **B,** UMAP plots show the expression of acknowledged markers of epithelial tumor, stromal and immune cells. Cells (dots) are colored by the expression level of each marker. Red indicates higher level while grey indicates lower level of expression.

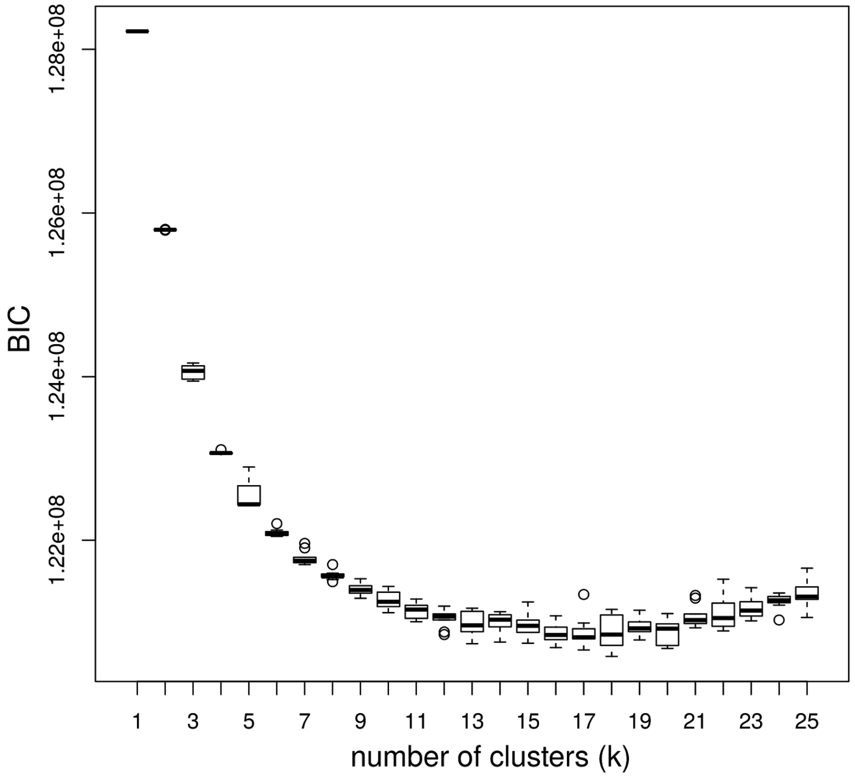


**Supplementary Figure S2**. **Selection of number of clusters based on Bayesian information criterion (BIC).** The model was fitted for $k = 1, 2, ..., 25$ with 10 different random initial parameter sets for each $k$, and $k = 12$ was selected based on BIC.

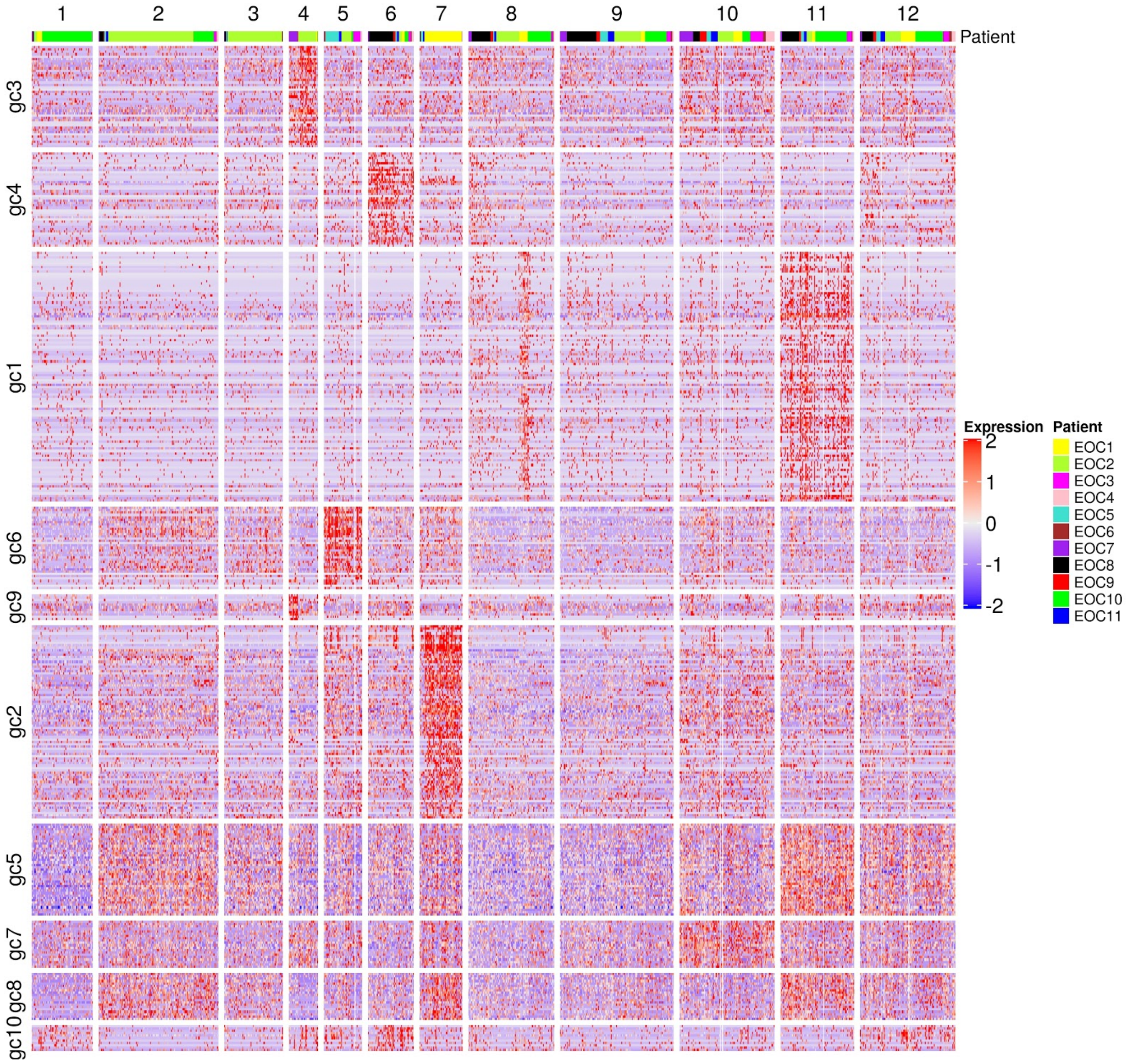


**Supplementary Figure S3**. Heatmap of the expression of the 10 distinct gene communities in the clusters detected from an alternative fitting with 200 random restarts. Rows correspond to genes and columns to cells. The color bar on the top denotes the patients.

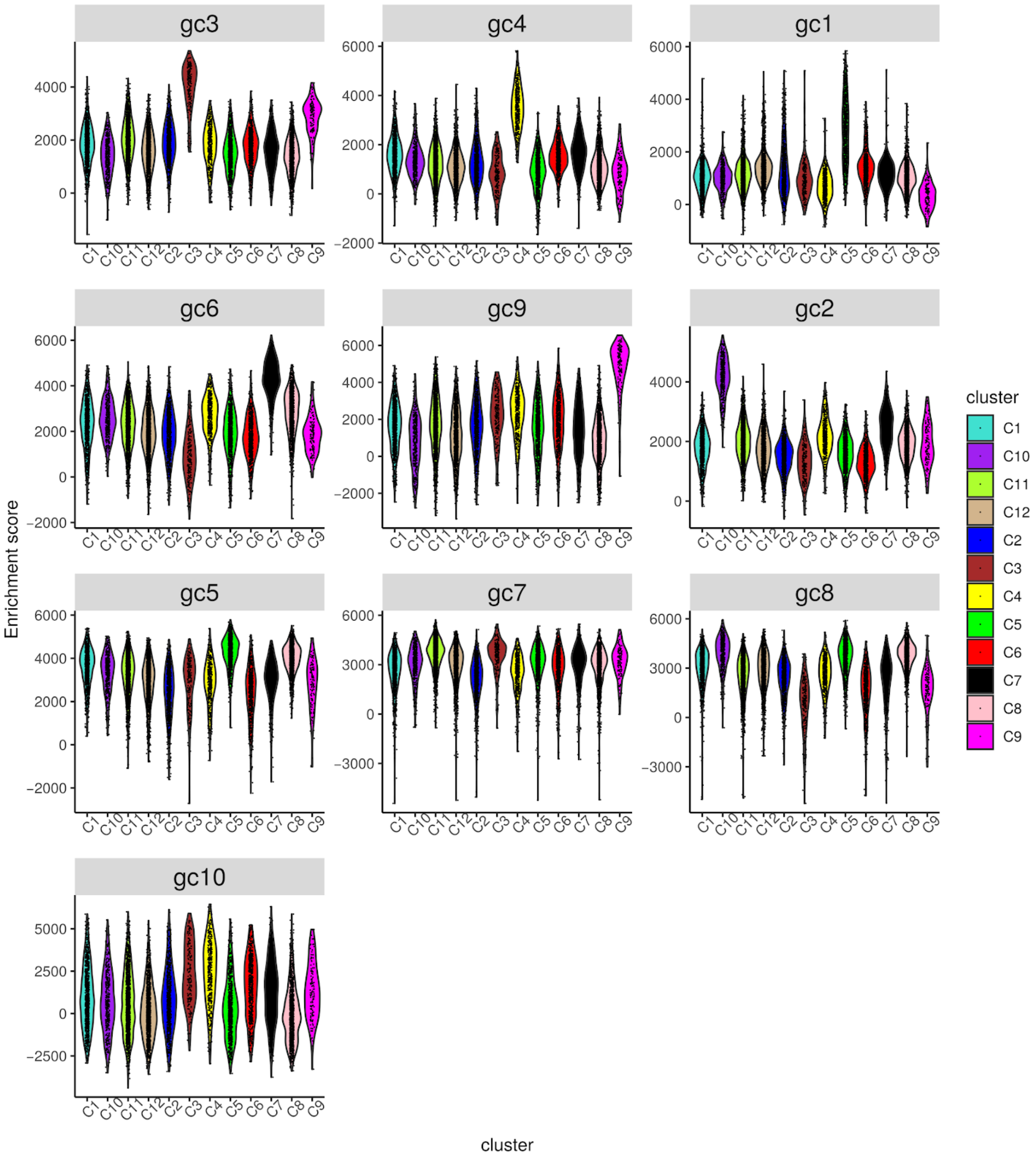


**Supplementary Figure S4**. Violin plots show single sample gene set enrichment (ssGSEA) scores of each gene community for each cell in each cell cluster. Colors denote cell clusters.

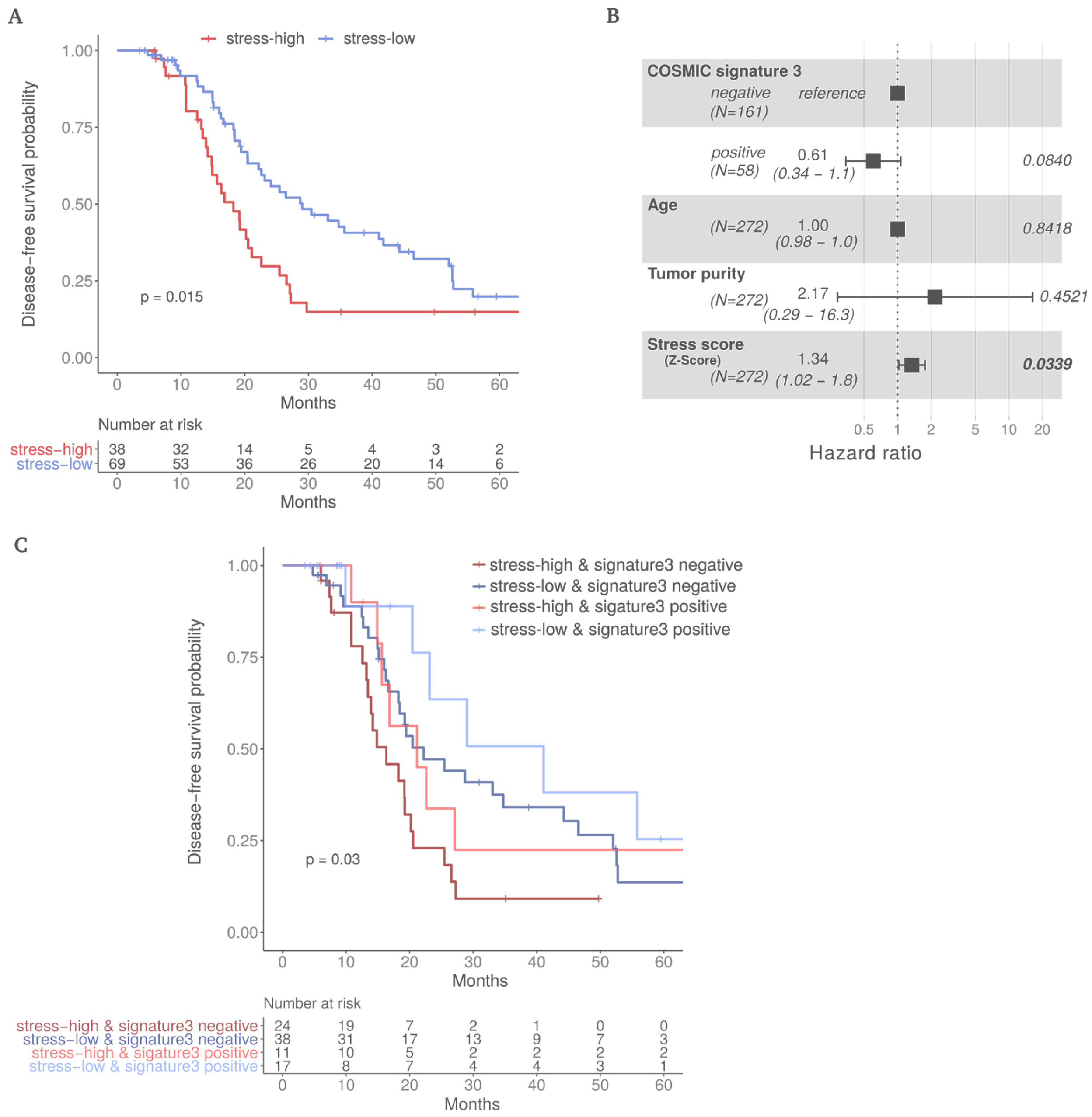


**Supplementary Figure S5. High stress score is associated with poor disease-free interval. A,** Kaplan–Meier curves on disease-free survival for stress-high and stress-low patients (log-rank test, *p* = 0.015) from the TCGA cohort. **B,** Forest plot showing hazard ratios, their confidence intervals and p-values based on a multivariate Cox proportional hazards regression model studying disease-free interval in relation to COSMIC signature 3 status, age at diagnosis, tumor purity and stress score. Significant p values (P < 0.05) are shown in bold. **C,** Kaplan–Meier curves on disease-free survival for patients from the TCGA cohort with stress-high & signature 3 negative (ii) stress-low & signature 3 negative (iii) stress-high & signature

3 positive and (iv) stress-low & signature 3 positive. The log-rank test p value is 0.03. The number of patients at risk is listed below the survival curves for each time point.

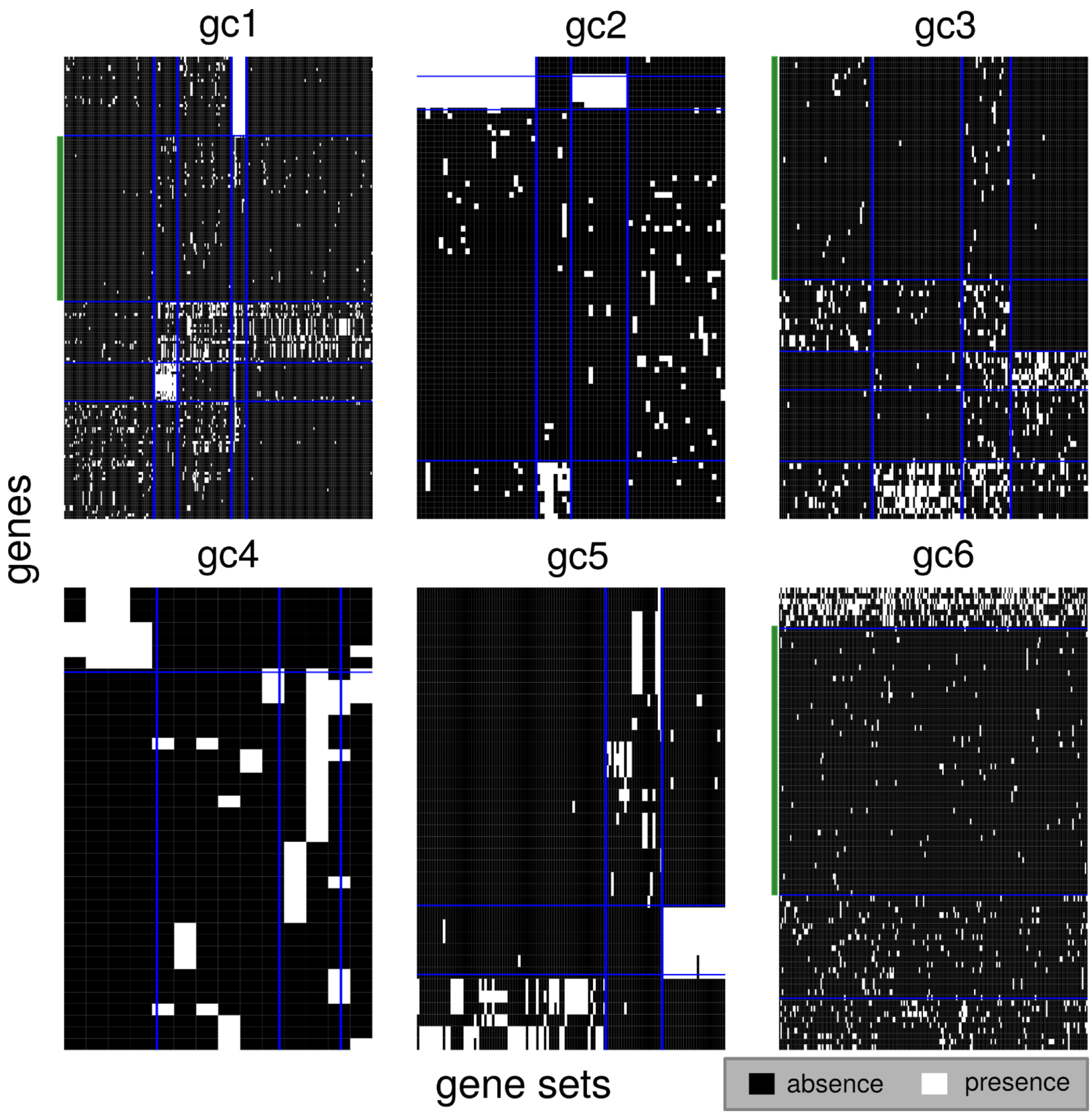


**Supplementary Figure S6**. **Filtering of gene communities based on functional annotation.** Biclustering was applied to the binary matrix of the presence/absence of each gene in each significantly overrepresented gene set. The heatmaps show the clustered binary matrix with blue grids indicating distinct clusters. Green bars on the left indicate the gene clusters that have less than 3% presence in any of the gene-set clusters, which were excluded from subsequent analysis.

**Supplementary Table S1**. List of genes in each gene community (Continued on the following page).

| Gene community | Genes |
| --- | --- |
| **gc1** | AURKA, AURKB, BIRC5, BUB1, BUB1B, CCNA2, CCNB1, CCNB2, CDC20, CDC25C, CDC45, CDC6, CDCA8, CDK1, CDT1, CENPA, CENPE, CENPF, CENPH, CENPK, CENPM, CENPN, CENPU, CKAP5, DHFR, E2F1, H2AFV, H2AFZ, HAUS1, HIST1H2BH, HIST1H3G,Data HIST1H4C, HIST2H2AC, KIF18A, KIF20A, KIF23, KIF2C, LIG1, LMNB1, MAD2L1, MCM10, MCM2, MCM3, MCM4, MCM5, MCM6, MCM7, MCM8, NDC80, NEK2, NUF2, ORC6, PCNA, PLK1, PLK4, POLD1, POLD2, PRIM1, PTTG1, RFC2, RFC3, RFC4, RRM2, SGO1, SGO2, SKA1, SKA2, SKP2, SPC24, SPC25, TK1, TMPO, TPX2, UBE2C, VRK1, ZWILCH, ZWINT, BRCA1, CHEK1, CLSPN, RHNO1, RMI2, FANCG, HMGB2, KIF4A, RRM1, TYMS, RACGAP1, CBX5, BRCA2, FANCD2, RAD51, RAD51AP1, FANCB, FANCI, UBE2T, KIF11, KIF15, KIF20B, KIF22, KIFC1, HIST1H1A, HIST1H1B, GGH, DTYMK, AKR1B1 |
| **gc2** | CTSL, HLA-DMA, HLA-DMB, HLA-DOA, HLA-DPA1, HLA-DPB1, HLA-DQA1, HLA-DQA2, HLA-DRA, HLA-DRB5, HSP90AB1, HSPA2, HSPA4, HSPA5, HSPA6, PDIA3, CPE, CTSF, C1R, CXADR, SFTPD, TUBB2A, TUBB6, FOLR1, GAS6, NPC2, RUNX1, TGFBR2, GNAI1, RBPJ, IL11RA, HIST1H2AE, DNAJC3, FBXO2, HERPUD1, HSP90B1, SKP1, UBE2D1, UBQLN1, NECTIN2, SDC2, CCND3, DNAJB9, ERP27, SIAH1, PPP2CA, BAG3, HSPA12A, BMP2, LATS2, PPP2R2B, PRKCI, WNT6, IFITM1, IFITM2, CRMP1, DPYSL3, PLXNA2, MYL9, TNNC1, TNNI1, TNNT2, TLE1, TLE4, APP, PELI2, VAMP2, DNAJA4, PLAT, ACADVL, TSPYL2, NR1D2, PURA, GATA6, TWIST1, RING1, BAMBI, ATG101, RAPGEF2, NR4A3, SOX4, SLC22A18 |
| **gc3** | DST, EGFR, EPHB2, ITGA6, PIK3CD, PLEC, SFN, SMAD3, CLTC, EIF2AK2, ITGA2, KPNA1, STAT1, BCAR1, CCND1, COL1A2, ITCH, MMP1, MMP12, PML, TNC, CDK6, FADD, FAS, MX1, E2F3, BIRC2, CAPN2, VASP, MMP9, DFFA, KPNB1, ADAMTS5, MMP10, AP2B1, ASAP1, SDCBP, CYP1A1, CYP1B1, CYP3A5, MAFG, SLC7A11, IGF2 |
| **gc4** | B3GNT7, B3GNT8, GCNT3, MUC15, MUC4, MUC5B, ST3GAL4, LCN2, S100A8, S100A9, ALOX5, AOC1, B2M, C3, CEACAM1, CEACAM6, MGST1, OSTF1, PPBP, S100P, SLPI, TMEM173, DSC2, EVPL, IVL, KRT13, KRT4, KRT6A, KRT6C, HRASLS2, LPCAT4, PLBD1, RARRES3, CDKN2B, STEAP4, TGFB1, CFB, CCL28, CX3CL1, CXCL10 |

| | |
|---|---|
| **gc5** | AIMP1, EIF3I, MRPL13, MRPL15, MRPL16, MRPL2, MRPL28, MRPL3, MRPS7, RPN2, SRP9, TSFM, PSMA4, PSMB5, PSMB6, PSMB7, PSMC5, PSMD8, ACTL6A, BANF1, PPIA, XRCC6, VDAC3, FH, MDH2, SDHB, TPI1, ECHS1, HSD17B10, POLR2G, NDUFB6, HNRNPA2B1, HNRNPA3, SNRPD3, PRDX3, PTMA, CCT7, VBP1, COPE |
| **gc6** | CEBPB, CEBPD, FOS, IL6, JUN, JUNB, MCL1, MYC, SOCS3, ATF3, DUSP1, EGR1, FOSB, CEBPA, DDIT3, EGR2, CDKN1A, GADD45B, TNF, HES1, HBEGF, BCL6, NR4A1, DUSP6, GADD45G, ID2, NFKBIA, PLK3, SNAI2, CREB5, HLA-G, HIST1H2BC, HIST1H2BG, CALML3, SNAI1 |
| **gc7** | BMS1, DCAF13, MPHOSPH10, PNO1, ACIN1, ROCK1, TJP1, ESCO1, WAPL, EIF1AX, PNN, RNPS1, DYNC1LI1, BDP1, CRCP, PAFAH1B1, PRPF40A, SFSWAP, KMT2A, SETD2 |
| **gc8** | GLO1, IDH3B, NDUFA10, NDUFB5, PDHA1, SUCLG1, TRAP1, UQCRC1, UQCRC2, MDH1, ALDH7A1, DECR1, ECH1, ECI2, PCCB, SHMT1, HACD3, PTGR1, MGST2, PRDX6 |
| **gc9** | CCL20, CXCL1, IL1R2, IL1RN, SEC11C, SPCS3, BIK, BIRC3, CDKN2A, DAB2, GJB2 |
| **gc10** | GBP4, IFI27, IFI35, IFIT1, ISG15, OAS1, OAS2, STAT2, TRIM22, TAP1, ITGAV |

**Supplementary Table S2**. Summary of the functional enrichment of each gene community (Continued on the following page).

| cell cluster | gene community | biological process |
|---|---|---|
| C3 | gc3 | TGF-beta Signaling Pathway, Focal Adhesion |
| C4 | gc4 | O-linked glycosylation of mucins |
| C5 | gc1 | Cell Cycle, DNA repair |
| C7 | gc6 | (Stress related) ILs-mediated signaling events, Adipogenesis, Apoptosis, Cellular responses to stress |

| | | |
|---|---|---|
| C9 | gc9 | Interleukin-10 signaling |
| C10 | gc2 | Antigen processing and presentation, MHC class II antigen presentation |
| C3, C4 | gc10 | Interferon Signaling |
| C3, C11 | gc7 | rRNA processing, Apoptosis |
| C5, C8 | gc5 | Proteasome |
| C5, C8, C10 | gc8 | TCA cycle |
